# Indigenous Cultural Safety Training for Applied Health, Social Work and Education Professionals: A PRISMA Scoping Review

**DOI:** 10.1101/2022.10.06.511097

**Authors:** Tammy MacLean, Jinfan Qiang, Lynn Henderson, Andrea Bowra, Lisa Howard, Victoria Pringle, Tenzin Butsang, Emma Rice, Erica Di Ruggiero, Angela Mashford-Pringle

## Abstract

**Background:** Anti-Indigenous racism is a widespread social problem in health, social work, and education systems in English-speaking Colonized countries such as Canada, with profound negative impacts to the health and education of Indigenous peoples. In 2015, Canada’s Truth and Reconciliation Commission recognized the legacy and impact of Colonization and recommended training programs for these professions on cultural competency and curricula, and on the colonial history of Canada. Yet there is little evidence on best practices for such training, highlighting the need to synthesize existing findings on how these training programs are developed, implemented, and evaluated.

**Methods:** This scoping review explored the academic literature on Indigenous cultural safety and competence training in the health, social work, and education fields. Medline, EMBASE, CINAHL, ERIC and ASSIA were searched for articles published between 1996-2020 in Canada, United States, Australia, and New Zealand. The Joanna Briggs Institute’s three-step search strategy was used as was the PRISMA extension for Scoping Reviews. Data was charted and synthesized in three stages.

**Results:** 134 were included in this review. Data was extracted on four themes: 1) Article Characteristic; 2) Cultural Safety Concepts, Critiques and Rationale; 3) Characteristics of Cultural Safety Training; and 4) Evaluation Details of Cultural Safety Training. Findings suggest that research on cultural safety training programs in health, social work and education has grown significantly. Nursing and medicine professions have received a significant proportion of cultural training programs, compared with general/allied health, social work, and education. Across fields, professionals and students were targeted equally by training programs. Only half of evaluations of cultural safety and related intervention identified methodological limitations.

**Implications:** Considering, comparing, and contrasting literature on cultural safety and related concepts and how they are applied in practice would advance this scholarly work, as would more robust evaluations of cultural safety and similar training interventions to understand their impact at the individual level. Finally, commitment to meaningfully engage Indigenous communities to develop, implement and evaluate such programs is urgently needed.

## 1. INTRODUCTION

### Rationale

There is substantial evidence to suggest that anti-Indigenous racism is a widespread health and social problem in Canada. While the term, Indigenous, refers to a variety of Aboriginal groups around the world (1), in Canada, Indigenous refers to the country’s first inhabitants, namely First Nations, Inuit, and Métis peoples, as established in Section 35 of the Canadian Constitution (2). Racism may be understood as “racist ideologies, prejudiced attitudes, discriminatory behaviour, structural arrangements and institutionalized practices resulting in racial inequality as well as the fallacious notion that discriminatory relations between groups are morally and scientifically justifiable” (3). According to several population-based studies undertaken with Indigenous peoples across Canada, between 39% and 43% of respondents reported experienced racism (4-7). Systemic anti-Indigenous racism in health service organizations across the country is equally problematic. The investigations into the deaths of Brian Sinclair in Manitoba in 2008 (8) and Joyce Echaquan in Quebec in 2020 (9), coupled with the alarming findings from the anti-Indigenous racism investigation in British Columbia’s health care system in 2020 (10) collectively demonstrate the insidious nature of anti-Indigenous racism in Canada’s health systems. Not only is racism a serious challenge for Indigenous peoples while accessing care, but also it has profound negative impacts on their likelihood of accessing future care (10-17).

Beyond the health sector, research evidence also points to widespread anti-Indigenous racism in Canada’s social work (18, 19) and Kindergarten to Grade 12 education systems (20, 21). Moreover, Indigenous peoples have been identified as the most disadvantaged with respect to accessing education in English speaking, colonized countries such as Canada, United States, Australia and New Zealand (20). Collectively, this evidence runs contrary to the inalienable rights established and set out within the United Nations Declaration on the Rights of Indigenous Peoples, including:

> the right to be actively involved in developing and determining health […] and social programmes affecting them […] the right to access, without any discrimination, all social and health services […], the right to enjoyment of the highest attainable standard of physical and mental health […], [and] the right to all levels and forms of education [in the public system] […] and that indigenous peoples, in the exercise of their rights, should be free from discrimination of any kind. (22)

In 2015, Canada’s Truth and Reconciliation Commission (TRC) recognized the legacy and impact of residential schools on Indigenous peoples across the country (23) and put forth 94 Calls to Action, including for the health, social work/child welfare, and education systems. These calls included providing cultural competency training for health professionals to address unconscious bias and systemic racism and developing culturally appropriate education curricula, while building student capacity for intercultural understanding, empathy, and mutual respect (24). The calls also set out the need to ensure that social workers who conduct child-welfare investigations to be properly educated and trained about the history and impacts of residential schools on children and their caregivers. In terms of education systems, the TRC recommended that resources be provided to ensure Indigenous schools utilize Indigenous knowledge and teaching methods in the classroom and that relevant teacher-training needs were identified to ensure such knowledge and methods were implemented.

Initiatives such as Canada’s TRC, along with similar federal government initiatives in Australia and New Zealand have led to the emergence of new training programs to facilitate cultural safety and competence, and ultimately, to eliminate anti-Indigenous racism and discrimination in health, social work and education systems (25, 26). However, given the nascency of the field of Indigenous cultural safety training combined with the need to expand the availability of such training to address anti-Indigenous racism, there is an urgent need to synthesize and understand existing evidence on how such training programs are conceptualized and developed, as well as how they are implemented in practice and evaluated for impact.

### 1.2 Objectives

Our team sought to explore the academic literature that conceptualizes and/or operationalizes Indigenous cultural safety training within the fields of health, social work, and education. The aim of this study was to answer the following three questions: 1) What is the general state of knowledge on Indigenous cultural safety training in the applied fields of health, social work, and education? 2) What methods are used to develop, implement, and evaluate Indigenous cultural safety training for students and professionals in the applied fields of health, social work, and education? And 3) What content is included in existing Indigenous cultural safety training program for students and professionals in the fields of health, social work, and education?

## 2. METHODS

### 2.1 Protocol and registration

A detailed research protocol for this scoping review is published in *Social Science Protocols* doi.org/10.7565/ssp.2020.2815 (27), an open-access online journal platform. The authors of this paper chose not to register this study, both because of the rigid methodological requirements for registration, which do not align well with Indigenous research approaches, and because no other Indigenous studies were registered at the time. Indigenous cultural safety is a nascent area of research and practice, and the goal of this review was to achieve a comprehensive search. Our analysis sought to draw out insights that could inform Indigenous cultural safety training for professionals in the applied fields of health, social work, and education, which work with Indigenous peoples in Canada.

### 2.2 Eligibility Criteria

We reviewed the global academic literature on Indigenous cultural safety training that included but was not limited to Indigenous peoples in Canada. Inclusion criteria: This review was restricted to articles about Indigenous cultural safety in health, social work and education in British colonial settler nation states, including Australia, New Zealand, Canada, and the United States. All peer-reviewed primary research articles on the topic of Indigenous cultural safety within the fields of health, social work, and education that were reported in the academic literature between 1996-2020 were included. The review dates were selected to ensure that the full history of the term “cultural safety”, first coined in 1996 (28), was captured. This review included all articles published in or translated into English. Exclusion criteria

### 2.3 Information Sources

To identify potentially relevant literature in health, social work, and education, the following five bibliographic databases were searched from 1996-2020: Medline, EMBASE, CINAHL, ERIC and ASSIA. These databases were selected to capture the fields of health, education, and social work as aligned with the focus of this review. The search strategies were developed by an experienced librarian from Gerstein Library at the University of Toronto and further refined through team discussion.

### 2.4 Search Strategy

A three-step search strategy (Aromatis & Munn, 2020) was utilized for this review. The first step involved a limited search of two initial databases, Medline and EMBASE, followed by an analysis of subject headings and search terms based on identified titles and abstracts. Table 1 (Search Strategy for MEDLINE AND EMBASE (Ovid)) outlines the keyword searches employed at this step. A second search was then conducted using all identified subject headings and keywords across all five databases. Table 2 and Table 3 detail the keyword searches employed to search CINAHL, and ERIC and ASSIA respectively (for details of this process, see section 2.5: Selection of Sources of Evidence). These searches were conducted in May 2020, and collectively resulted in 3600 studies, which were imported into Covidence for screening. The decision was made by our team to only search the literature prior to COVID-19.

**Table 1:**
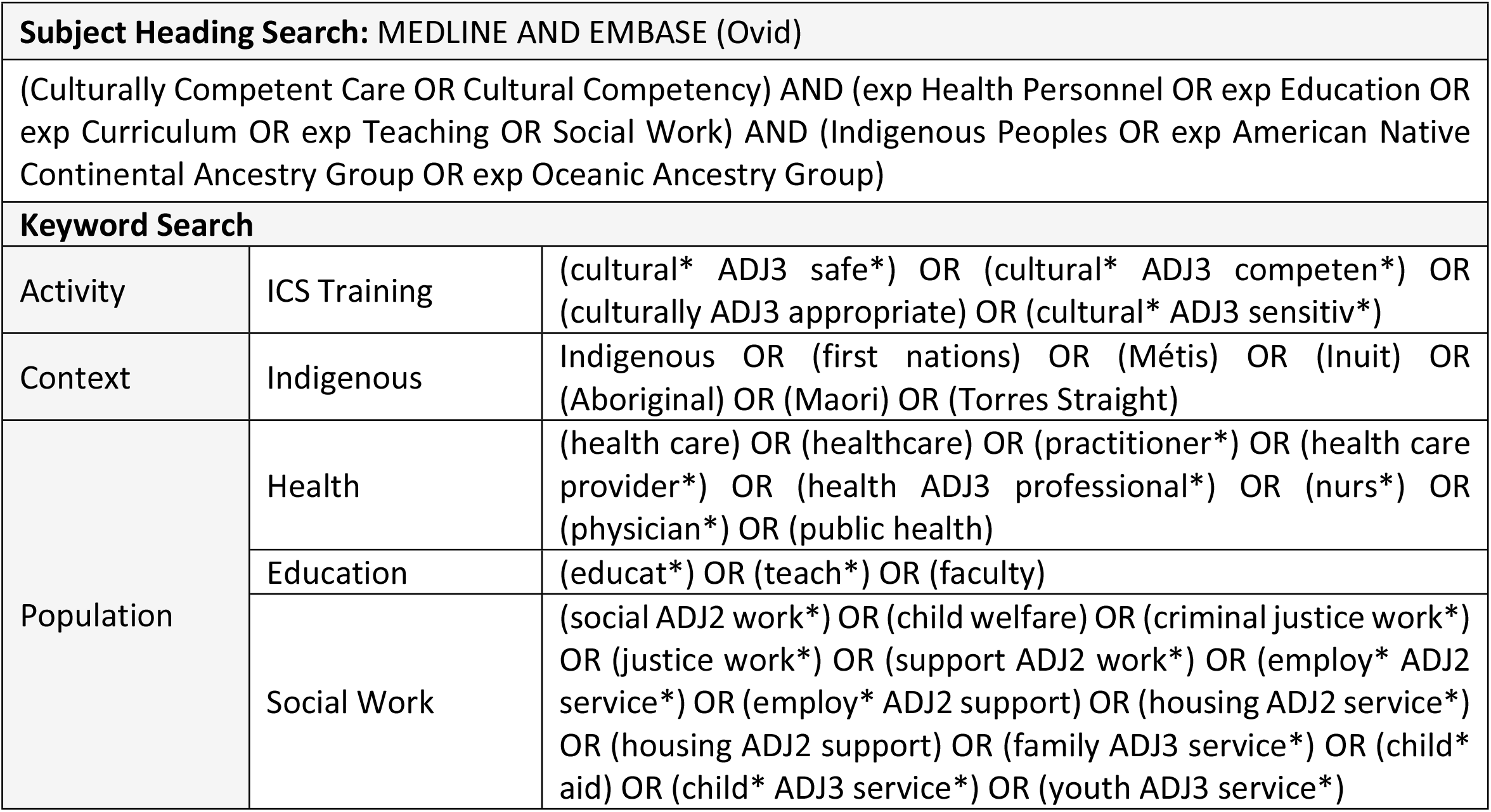
Search Strategy for MEDLINE AND EMBASE (Ovid)

| **Subject Heading Search: MEDLINE AND EMBASE (Ovid)** |  |  |
| --- | --- | --- |
| (Culturally Competent Care OR Cultural Competency) AND (exp Health Personnel OR exp Education OR exp Curriculum OR exp Teaching OR Social Work) AND (Indigenous Peoples OR exp American Native Continental Ancestry Group OR exp Oceanic Ancestry Group) |  |  |
| **Keyword Search** |  |  |
| Activity | ICS Training | (cultural* ADJ3 safe*) OR (cultural* ADJ3 competen*) OR (culturally ADJ3 appropriate) OR (cultural* ADJ3 sensitiv*) |
| Context | Indigenous | Indigenous OR (first nations) OR (Métis) OR (Inuit) OR (Aboriginal) OR (Maori) OR (Torres Straight) |
| Population | Health | (health care) OR (healthcare) OR (practitioner*) OR (health care provider*) OR (health ADJ3 professional*) OR (nurs*) OR (physician*) OR (public health) |
|  | Education | (educat*) OR (teach*) OR (faculty) |
|  | Social Work | (social ADJ2 work*) OR (child welfare) OR (criminal justice work*) OR (justice work*) OR (support ADJ2 work*) OR (employ* ADJ2 service*) OR (employ* ADJ2 support) OR (housing ADJ2 service*) OR (housing ADJ2 support) OR (family ADJ3 service*) OR (child* aid) OR (child* ADJ3 service*) OR (youth ADJ3 service*) |

**Table 2:** Search Strategy for CINAHL

| **Subject Heading Search** |  |  |
| --- | --- | --- |
| (Cultural Safety) OR (Cultural Competence) OR (Cultural Sensitivity) AND (Indigenous Peoples) AND (Social Work) OR (Social Work Service) OR (Students, Social Work) OR (Education, Social Work) OR (Social Service Assessment) OR (Education) OR (Health Personnel) OR (Facilities, Manpower and Services) OR (Occupational Health Services) |  |  |
| **Keyword Search** |  |  |
| Activity | ICS Training | (cultural* N3 safe*) OR (cultural* N3 competen*) OR (culturally N3 appropriate) OR (cultural* N3 sensitiv*) |
| Context | Indigenous | Indigenous OR (first nations) OR (Métis) OR (Inuit) OR (Aboriginal) OR (Maori) OR (Torres Straight) |
| Population | Health | (health care) OR (healthcare) OR (practitioner*) OR (health care provider*) OR (health N3 professional*) OR (nurs*) OR (physician*) OR (public health) |
|  | Education | (educat*) OR (teach*) OR (faculty) OR (curriculum) |
|  | Social Work | (social N2 work*) OR (child welfare) OR (criminal justice work*) OR (justice work*) OR (support N2 work*) OR (employ* N2 service*) OR (employ* N2 support) OR (housing N2 service*) OR (housing N2 support) OR (family N3 service*) OR (child* aid) OR (child* N3 service*) OR (youth N3 service*) |

**Table 3:** Search Strategy for ERIC and ASSIA (Proquest)

| Keyword Search |  |  |
| --- | --- | --- |
| Activity | ICS Training | (cultural* NEAR/3 safe*) OR (cultural* NEAR/3 competen*) OR (culturally NEAR/3 appropriate) OR (cultural* NEAR/3 sensitiv*) |
| Context | Indigenous | Indigenous OR (first nations) OR (Métis) OR (Inuit) OR (Aboriginal) OR (Maori) OR (Torres Straight) |
| Population | Health | (health care) OR (healthcare) OR (practitioner*) OR (health care provider*) OR (health NEAR/3 professional*) OR (nurs*) OR (physician*) OR (public health) |
|  | Education | (educat*) OR (teach*) OR (faculty) |
|  | Social Work | (social NEAR/2 work*) OR (child welfare) OR (criminal justice work*) OR (justice work*) OR (support NEAR/2 work*) OR (employ* NEAR/2 service*) OR (employ* NEAR/2 support) OR (housing NEAR/2 service*) OR (housing NEAR/2 support) OR (family NEAR/3 service*) OR (child* aid) OR (child* NEAR/3 service*) OR (youth NEAR/3 service*) |

### 2.5 Selection of Sources of Evidence

The selection of studies for this review was performed in Covidence independently by two reviewers and involved a four-stage process (see Image 1: Prisma Flow Diagram, in Appendix). The first stage of screening involved identifying and removing 1117 duplicate studies from Covidence (see results in Table 4: Scoping Search Results). The second stage of screening involved reviewing the titles and abstracts of the 2483 remaining article and applying the inclusion and exclusion criteria chosen for the study. The inclusions criteria include the following: 1) article focuses on an Indigenous cultural safety training program (as defined above) for students/professionals in the fields of health, education, or social work; 2) article pertains to investigations conducted in the English speaking, colonial settler countries of Australia, New Zealand, Canada, or the United States; 3) published as peer reviewed articles; 4) published in English; and 5) published since 1996. The exclusion criteria for the first stage of screening included: 1) describes a training program other than Indigenous cultural safety; 2) pertains to fields other than health, education, or social work; 3) published as literature reviews or conference summaries; 4) published prior to 1996; and 5) published in a language other than English. Applying the exclusion criteria resulted in removing 2163 articles from the study.

**Table 4:** Scoping Search Results

| Source | Studies Imported | Duplicates Removed | Studies Selected |
| --- | --- | --- | --- |
| Ovid (MEDLINE, EMBASE) | 1,372 | 5 | 1,366 |
| CINAHL | 1,608 | 820 | 788 |
| Proquest (ERIC, ASSIA) | 621 | 292 | 329 |
| Additional articles retrieved from reviewing study reference lists | 0 | 0 | 0 |
| **Total** | **3,600** | **1,117** | **2,483** |

The third stage of screening involved reviewing the full text articles to the remaining 320 titles and applying the same inclusion and exclusion criteria described in the previous screening stage. This process resulted in the exclusion of 178 additional studies for the following reasons: not related to cultural safety training and/or cultural safety concepts (n=126); not peer-reviewed (n=36); conference summaries (n=8); duplicate content (n=2); literature reviews; (n=2) not focused on targeted professional groups (n=2); full text not found (n=1); published before 1996 (n=1). As the removal of these articles, 142 titles were selected for extraction. The reference lists of these remaining articles were then searched for additional studies that met the inclusion and exclusion criteria, and no further studies were identified during this process. Finally, an additional 3 articles were excluded during the extraction process for containing duplicate content to that contained in studies already selected (n=2) and because a full text document could not be located (n=1). A total of 134 articles were extracted for this study.

### 2.6 Data Charting Process

A data-charting tool was developed for this study by AB with guidance from co-authors Dr. Mashford-Pringle, Dr. Erica Di Ruggiero, and a Librarian from Gerstein library, University of Toronto. The data-charting tool was developed in Google Spreadsheet by the first author of the published protocol paper for the study (27). While developing the data charting tool, several decisions were made to establish the focus, nature, and scope of data to be extracted. These decisions were: 1) to cut and paste entire ‘chunks’ of text verbatim into the tool rather than paraphrasing the data; 2) to include article page numbers associated with data on cultural safety concepts and critiques; and 3) to include both the study aim, along with the research question if explicitly stated and relevant to the study. The data-charting tool was then piloted by all members of the review team based on two pre-selected articles. Several procedural decisions were made during the pilot stage to further articulate the scope of data to chart, both for existing data points and for two new data points added during this stage. These decisions included: 1) whether the paper included Indigenous authors or Elders; 2) identifying whether a data collection or evaluation tool is included or referenced; 3) specifying that ‘training modality’ refers to the format (e.g. in-person, online etc.) and ‘training components’ relate to the topics covered; 4) including only ‘yes’ or ‘no’ on four specific data points; 5) specify that ‘duration of time the training program has been running’ is in relation to the study publication year; 6) adding ‘recommendations outlined in the article’; and 6) adding ‘stated limitations of the evaluation’.

The data-charting process involved charting each article twice and this was undertaken between August 2020 and June 2021. The first author (TM) independently completed one round of data-charting and the second round was completed by a team of six reviewers independently, with a large majority of the second extraction (102/134 articles) carried out by three of the six reviewers (LA, RQ and LH). The decision to involve six reviewers was made to expedite the extraction process, as the study results were needed to inform the development of pilot Indigenous cultural safety training intervention for Faculty members, staff and students of the Faculties of Nursing, Medicine, Social Work and Public Health at the University of Toronto. Several questions arose among the reviewers throughout the charting process concerning the nature and amount of the data to be extracted and charted, and these issues were generally resolved through discussion as a team. Any disagreements between two reviewers on a given paper were resolved through discussion or further adjudication by a third reviewer/co-author where necessary.

### 2.7 Data Items

The items included in the data-charting tool were developed deductively and addressed four main themes (see Table 5: Data Themes and Focus). The first theme includes *Article Characteristic* for Indigenous cultural safety training for the sources of evidence included in the review, namely: bibliographic details; country of study; whether author(s) identified as Indigenous; study aim/objective(s); target field(s)/discipline(s) (i.e. medicine/physicians; nursing/midwifery; allied health; health, general; education; and/or social work); and target population (students and/or professionals). The second theme concerns *Cultural Safety Concepts, Critiques and Rationale*, which includes cultural safety and related concepts either articulated or cited by study authors, along with critiques of the concepts, and the rationale given for why cultural safety is necessary in the fields of health, education, and social work.

**Table 5:** Data Themes and Focus

| Data Themes | Data Focus |
| --- | --- |
| **1. Article Characteristics** | Title<br>Journal<br>Publication Year<br>Country of Study<br>Authors<br>Did author(s) self-identify as Indigenous: [yes OR no]<br>Study aim/objective(s)<br>Target Field(s)/Discipline(s): [Medicine; Nursing/Midwifery; Allied Health; Health, General; Education; Social Work]<br>Target Population: [students; professionals] |
| **2. Cultural Safety Concepts, Critiques and Rationale** | Articulate/cite a cultural safety or related concept<br>Critique a cultural safety or related concept<br>Rationale for cultural safety |
| **3. Characteristics of Cultural Safety Training (if applicable)** | Did article describe a cultural safety training? [yes OR no]<br>[If yes, continue below]:<br>Who sponsored the training?<br>Who developed the training?<br>Is there evidence of Indigenous scholar, practitioner, knowledge keeper or community members in the development of the training [yes OR no]<br>Which Indigenous community partners were engaged?<br>Who delivered the training?<br>No. years training has run (r/t publication year)<br>Who received the training [Profession; Setting (rural OR urban)]?<br>Training objectives<br>Modalities used to deliver the training<br>Training duration/timeline<br>Brief description of training components<br>Number of people who completed the training<br>Details of post training supports provided<br>Stated limitations of the training |
| **4. Evaluation Details of Cultural Safety Training (if applicable)** | Did article present an evaluation of a cultural safety training? [yes OR no]<br>[If yes, continue below]:<br>Evaluation objective(s)<br>Evaluation data collection methods<br>Type of data collected (qualitative; quantitative)<br>Key evaluation results<br>Stated limitations of evaluation<br>Recommendations provided<br>Was a data collection tool provided [yes OR no] |

The third theme involves the *Characteristics of Cultural Safety Training* described in the sources of evidence. Data themes include who sponsored and developed the training program, whether Indigenous scholars, and practitioners or knowledge keepers were involved in the development process and the roles of community partners who were engaged. Other themes include who delivered and received the training program, along with details of the training program itself – such as the objectives, delivery modality, component descriptions, delivery duration and timeline, details of any post-program support, and the programs’ stated limitations. The fourth and final theme addresses *Evaluation Details of Cultural Safety Training*. This data involves the evaluation objectives, data collection methods, type(s) of data collected (i.e. qual, quant and both), evaluation results, stated evaluation limitations, recommendations for future cultural safety training programs, and whether a data collection tool was provided.

### 2.8 Synthesis of Results

The process of extracting data from selected articles and charting it into the data extraction tool was undertaken in one of two ways. For most data themes, the data was copied verbatim from selected studies and pasted into the data charting tool. However, there were four exceptions to this process, namely the following data themes that required only a yes/no response: 1) whether the author(s) self-identified as Indigenous; 2) whether there was evidence that Indigenous scholars, practitioners, knowledge keepers or community members were involved in developing the training; 3) whether the article describes an Indigenous cultural safety training; and 4) whether the study presented an evaluation of a cultural safety training. While the first two of these data themes made for useful data in and of themselves, the last two data themes were signposts to direct the reviewer to whether they should proceed and search within a given article to extract any data relevant to those themes.

Once the data was charted, the data synthesis process was undertaken in three stages and carried out by four reviewers (TM, RQ, LH, ER). The first stage involved analyzing both extractions of data for each theme to identify and resolve any discrepancies. All inconsistencies that were discovered were resolved through discussion and/or revisiting the sources of data where needed. The second stage involved analyzing data on the general characteristics of Indigenous cultural safety within the studies selected for the review, a process which involved categorizing the data within each data theme and then counting the frequency of findings within each category. The third stage involved synthesizing data that described a cultural safety training and/or the evaluation of a cultural safety training. The last stage involved synthesizing the data within each theme to create an overview of the data characteristics of Indigenous cultural safety, then categorizing the data within each theme into distinct groups, and finally, counting the frequency of findings within each category. Finally, the process of data synthesis also involved excluding or refocusing some of the data themes that were introduced in the protocol paper and proposed as part of this study. The reasons for these changes are varied and are set out in *Section 3.3: Evaluations of Indigenous Cultural Safety Training Interventions*.

## 3. RESULTS

### 3.1 Selection of Sources of Evidence

Based on the database searches detailed above, a total of 3,600 sources were imported into Covidence online screening (see Image 1: Prisma Flow Diagram, in Appendix). After removing 1117 duplicates and 2153 irrelevant studies, Covidence screened and assessed 320 full-text articles for eligibility. From these selected studies, and additional 178 were excluded during the analysis phase by our team for the following reasons: a) not related to cultural safety training and/or cultural safety conceptualization (n=126); b) not peer reviewed (n=36); c) conference summary (n=8); d) duplicate content (n=2); e) literature review (n=2); d) not focused on our target professional groups (n=2); e) full text of article not found/available (n=1); and f) published before 1996. While 142 remaining studies were selected for inclusion, an additional eight articles were removed during the process of data charting, because either the authors and content of these articles overlapped significantly with other selected articles, or the paper involved Indigenous peoples’ review of a cultural safety training but did not focus on the targeted professionals or students and their experiences with the training. Data from a total of 134 included full-text articles, including quantitative and/or qualitative research studies as well as expert opinion pieces, were extracted for this study.

### 3.2 General Characteristics of Sources of Evidence

Table 6 presents general characteristics from the 134 sources of evidence selected for this review. Over two-thirds (68%) of the studies were published between 2011-2020 (29-119), while over one-quarter (28%) were published between 2001-2010 (26, 120-155), and just over than 1 in 20 (4%) were published between 1996 and 2000 (28, 156-160). Approximately half of these studies (49%) were undertaken in Australia (36-39, 46, 47, 49, 53, 56, 58-60, 64, 66-69, 71, 72, 75, 77, 78, 80, 82, 83, 85-87, 89-91, 94, 95, 98, 99, 101-108, 110, 113, 115-119, 121-123, 126, 128, 130, 133, 136, 137, 140, 143, 144, 148, 151, 152, 160), with just over a fifth (21%) carried out in Canada (31-33, 35, 42-45, 50, 52, 55, 62, 63, 70, 74, 81, 84, 96, 100, 109, 111, 112, 114, 127, 132, 135, 145, 154). An estimated third of selected articles were either undertaken in the United States (16%) (30, 40, 41, 48, 61, 63, 65, 76, 79, 92, 93, 97, 120, 125, 134, 146, 153, 155, 156, 158, 159) or New Zealand (15%) (26, 28, 29, 34, 51, 54, 57, 73, 88, 124, 129, 131, 138, 139, 141, 142, 147, 149, 150, 157). One or more authors self-identified as Indigenous in only a fifth of the papers (20%) selected for this study (28, 30, 32, 34, 38, 41, 45, 47, 49, 55, 59, 62, 64, 66, 67, 71, 77, 79, 80, 83, 93, 104, 112, 118, 126, 136).

**Table 6.**
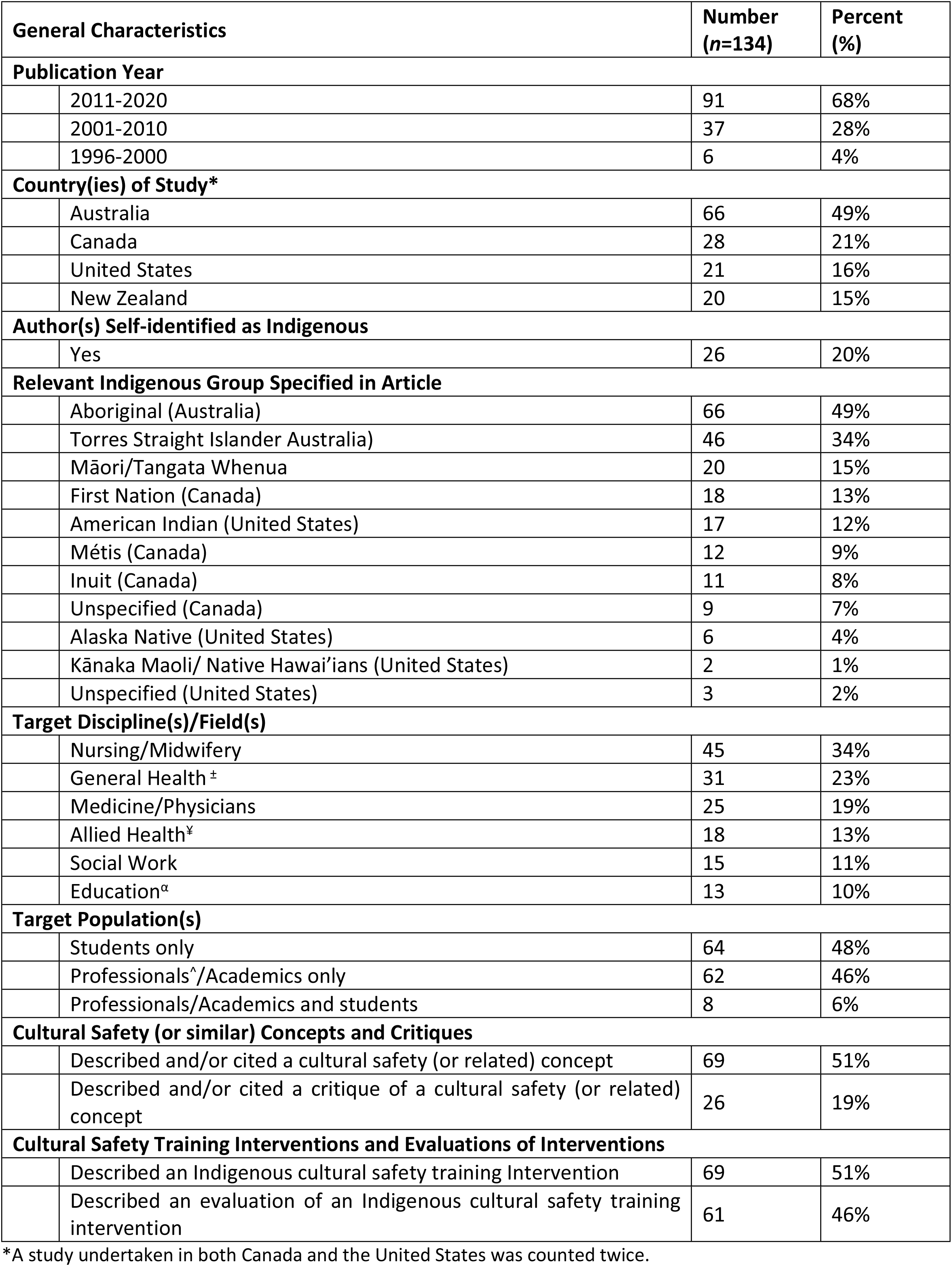
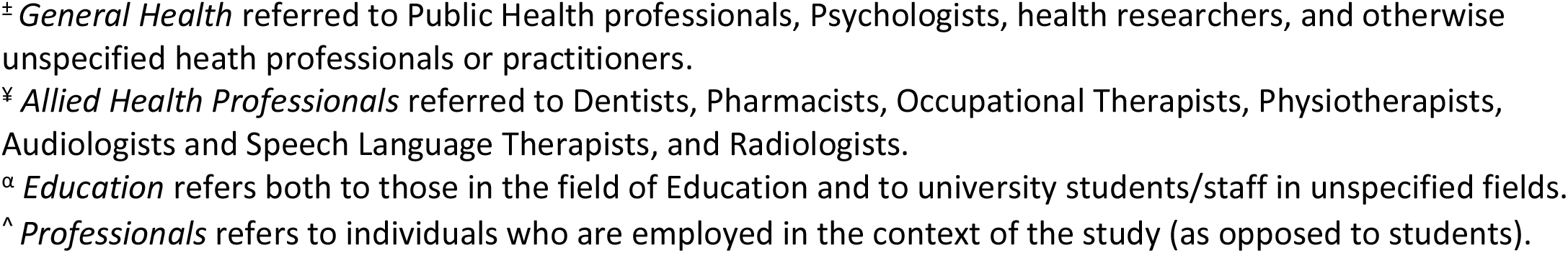
General Characteristics of Sources of Evidence

| General Characteristics |  | Number<br>(n=134) | Percent<br>(%) |
| --- | --- | --- | --- |
| **Publication Year** |  |  |  |
|  | 2011-2020 | 91 | 68% |
|  | 2001-2010 | 37 | 28% |
|  | 1996-2000 | 6 | 4% |
| **Country(ies) of Study\*** |  |  |  |
|  | Australia | 66 | 49% |
|  | Canada | 28 | 21% |
|  | United States | 21 | 16% |
|  | New Zealand | 20 | 15% |
| **Author(s) Self-identified as Indigenous** |  |  |  |
|  | Yes | 26 | 20% |
| **Relevant Indigenous Group Specified in Article** |  |  |  |
|  | Aboriginal (Australia) | 66 | 49% |
|  | Torres Strait Islander Australia) | 46 | 34% |
|  | Māori/Tangata Whenua | 20 | 15% |
|  | First Nation (Canada) | 18 | 13% |
|  | American Indian (United States) | 17 | 12% |
|  | Métis (Canada) | 12 | 9% |
|  | Inuit (Canada) | 11 | 8% |
|  | Unspecified (Canada) | 9 | 7% |
|  | Alaska Native (United States) | 6 | 4% |
|  | Kānaka Maoli/ Native Hawai'ians (United States) | 2 | 1% |
|  | Unspecified (United States) | 3 | 2% |
| **Target Discipline(s)/Field(s)** |  |  |  |
|  | Nursing/Midwifery | 45 | 34% |
|  | General Health<sup>±</sup> | 31 | 23% |
|  | Medicine/Physicians | 25 | 19% |
|  | Allied Health<sup>‡</sup> | 18 | 13% |
|  | Social Work | 15 | 11% |
|  | Education<sup>α</sup> | 13 | 10% |
| **Target Population(s)** |  |  |  |
|  | Students only | 64 | 48% |
|  | Professionals<sup>^</sup>/Academics only | 62 | 46% |
|  | Professionals/Academics and students | 8 | 6% |
| **Cultural Safety (or similar) Concepts and Critiques** |  |  |  |
|  | Described and/or cited a cultural safety (or related) concept | 69 | 51% |
|  | Described and/or cited a critique of a cultural safety (or related) concept | 26 | 19% |
| **Cultural Safety Training Interventions and Evaluations of Interventions** |  |  |  |
|  | Described an Indigenous cultural safety training Intervention | 69 | 51% |
|  | Described an evaluation of an Indigenous cultural safety training intervention | 61 | 46% |

<sup>‡</sup> *General Health* referred to Public Health professionals, Psychologists, health researchers, and otherwise unspecified health professionals or practitioners. <sup>¥</sup> *Allied Health Professionals* referred to Dentists, Pharmacists, Occupational Therapists, Physiotherapists, Audiologists and Speech Language Therapists, and Radiologists. <sup>α</sup> *Education* refers both to those in the field of Education and to university students/staff in unspecified fields. <sup>^</sup> *Professionals* refers to individuals who are employed in the context of the study (as opposed to students).

Among the Indigenous groups identified as relevant to selected articles, Aboriginal peoples from Australia comprised the greatest proportion (49%) (36-39, 46, 47, 49, 53, 56, 58-60, 64, 66-69, 71, 72, 75, 77, 78, 80, 82, 83, 85-87, 89-91, 94, 95, 98, 99, 101-108, 110, 113, 115-119, 121-123, 126, 128, 130, 133, 136, 137, 140, 143, 144, 148, 151, 152, 160), followed by Torres Straight Islanders also in Australia (34%) (36-39, 46, 47, 49, 53, 56, 60, 66-69, 71, 72, 80, 83, 85-87, 89, 98, 103-108, 110, 113, 115, 116, 118, 121-123, 126, 128, 130, 133, 137, 140, 148, 151, 152). Thereafter included Māori/Tangata Whenua in New Zealand (15%) (26, 28, 29, 34, 51, 54, 57, 73, 88, 124, 129, 131, 138, 139, 141, 142, 147, 149, 150, 157), First Nations in Canada (13%) and American Indians in the United States (12%) (30, 41, 48, 63, 65, 79, 92, 93, 97, 120, 134, 146, 153, 155, 156, 158, 159). Other Indigenous groups included Métis in Canada (9%) (31, 32, 44, 50, 52, 62, 63, 109, 112, 114, 135, 154), Inuit (8%) (31, 32, 44, 50, 52, 62, 63, 84, 109, 112, 135) and unspecified Indigenous peoples in Canada (7%) (43, 45, 55, 81, 96, 111, 127, 132, 145), with each identified in less than a tenth of articles. For publications from the United States, Alaska Natives (4%) (30, 61, 63, 93, 120, 159), Kānaka Maoli/Native Hawai’ians (1%) (76, 125), and unspecified Indigenous peoples (2%) (40, 74, 79) were mentioned in (or less than) one in twenty-five papers.

With respect to the target disciplines/fields within the articles, nursing/midwifery comprised the largest share at approximately one-third (34%) (26, 28, 39, 41, 42, 46-49, 51-55, 63, 72, 80, 85, 87-92, 106, 110, 111, 113, 121, 123, 124, 126-128, 130-133, 135, 136, 147, 149, 157, 158, 160). General Health Professionals – a category including public health professionals, psychologists, health researchers, and unspecified heath professionals or practitioners – were the target of nearly one-quarter (23%) of papers. Medicine (or physicians) was the target of nearly one-fifth of papers (19%) (26, 41, 43, 45, 47, 56, 57, 60, 62, 63, 76, 93, 94, 96, 108, 109, 111, 112, 114, 119, 121, 124, 125, 143, 150). Following the these three fields, allied health (13%), (26, 31, 32, 69-71, 73, 74, 81, 100, 101, 110, 121, 138, 140, 141, 144, 145), social work (11%) (58, 59, 64-68, 75, 76, 105, 111, 154-156, 159), and education (10%) (29, 33-35, 78, 82, 83, 102, 103, 118, 142, 146, 152), were the target fields for between one-seventh and one-tenth of papers. Nearly half of the studies targeted students of these disciplines (48%) (26, 28, 29, 32, 34, 39, 42-44, 46, 49, 52, 57, 59, 64-68, 70, 71, 73-78, 81, 83-87, 89, 91, 92, 95, 98, 99, 101-104, 106-109, 112-114, 118, 121, 123, 124, 135, 137, 139, 140, 142, 144, 150, 152, 153, 160), while a slightly smaller proportion targeted professionals and/or academics (46%) (30, 31, 35-38, 40, 41, 45, 47, 50, 51, 53-56, 58, 60-63, 69, 72, 79, 80, 82, 88, 94, 96, 97, 100, 105, 110, 111, 115-117, 120, 122, 126-133, 136, 138, 141, 143, 145-149, 154-Only one in twenty articles targeted both professionals/academics and students (6%) (33, 48, 90, 93, 119, 125, 134, 151).

In terms of the content within the articles selected for this review, just over half of the articles (51%) either articulated or cited a cultural safety (or similar) concept (e.g. cultural safety, cultural sensitivity, cultural awareness, cultural competence etc.) (28, 31-34, 38, 39, 44-47, 49-55, 59, 63, 64, 68, 71, 72, 79, 82, 84-86, 88-90, 92, 94, 96, 97, 100, 102, 106, 107, 109-111, 113, 114, 116, 121-123, 125, 126, 128-132, 136, 138, 139, 141, 144, 145, 147, 149-151, 154, 157, 159). Additionally, an estimated one-fifth of papers (19%) either articulated or cited a *critique* of a cultural safety or similar concept (28, 31, 38, 44, 50, 53-55, 59, 63, 79, 85, 97, 106, 107, 113, 122, 127, 128, 130-132, 147, 150, 151, 157). With respect to training interventions, approximately half of the selected articles (n=69; 51%) described a cultural safety training intervention (26, 32, 34-37, 39, 41-43, 47-49, 57, 58, 60, 62, 65, 66, 68-70, 72, 74-81, 83-86, 89, 91-93, 95, 101-108, 110, 111, 113, 114, 117, 119, 121, 123, 124, 134, 135, 137, 140, 142-144, 146, 148, 153, 154, While somewhat less than half of the papers (n=61; 46%) described an evaluation of a cultural safety training intervention (32, 34, 35, 39, 41, 43, 46, 47, 49, 51, 54, 56-58, 60-62, 64-66, 70, 72, 74, 76-78, 80, 81, 84, 86, 89, 91-93, 95, 98, 99, 101-108, 110, 111, 113, 117, 123, 124, 133, 134, 137, 144, 146, 148, 153, 154, 156, 158).

### 3.3 Details of Indigenous Cultural Safety Training Interventions

The details of training interventions for Indigenous cultural safety are presented in Table 7: Details of Indigenous Cultural Safety Training Interventions). Of these papers (n=69), more than two-thirds mentioned the involvement of Indigenous peoples in the training development process (n=48; 70%) (32, 34, 37, 39, 41-43, 46, 48, 49, 57, 58, 60, 62, 65, 68, 70, 75-79, 81, 83, 85, 86, 92, 93, 95, 101-104, 106-108, 110, 113, 114, 117, 123, 137, 140, 143, 144, 146, 148, 153). A wide variety of teaching approaches to delivering cultural safety content were illustrated as part of these interventions, with all having described more than one modality. Teaching/lectures was the most common training modality, described in almost a third of papers (n=21; 30%) (32, 34, 39, 41, 43, 46, 49, 57, 75, 76, 78, 83, 84, 86, 93, 95, 101, 113, 121, 124, 137). Workshops (n=18; 26%) (47, 57, 60, 62, 69, 72, 77, 79, 80, 93, 102, 103, 110, 117, 134, 137, 140, 148), discussions sessions (n=16; 23%) (32, 39, 41, 42, 47, 72, 76, 77, 85, 101, 102, 110, 111, 121, 135, 140), and immersive experiences/community visits (n=13; 19%) (32, 41, 43, 57, 76, 78, 92, 102, 106, 108, 140, 146, 153) were also common, detailed by estimated quarter to approximately a fifth of relevant articles.

**Table 7:**
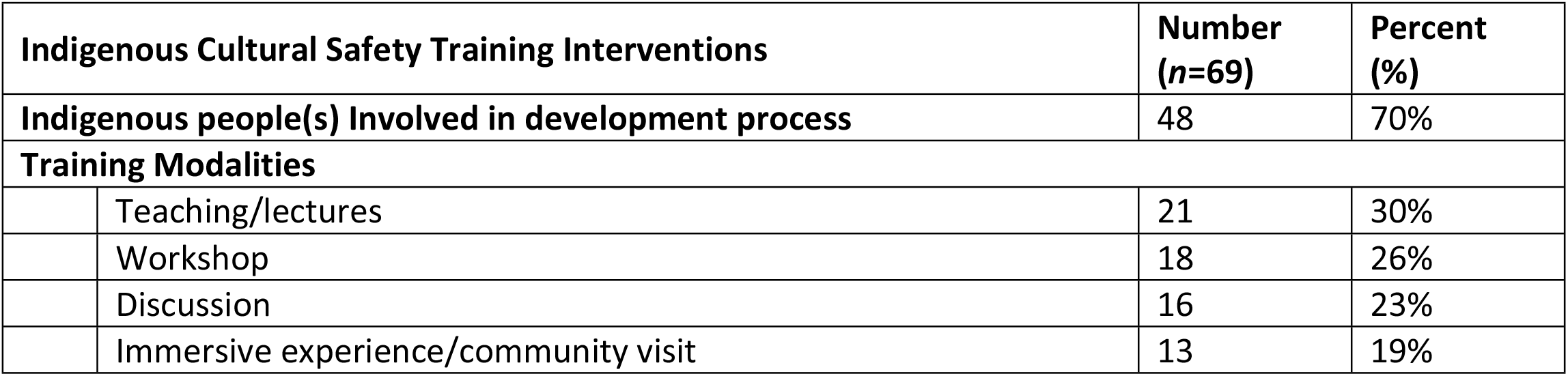
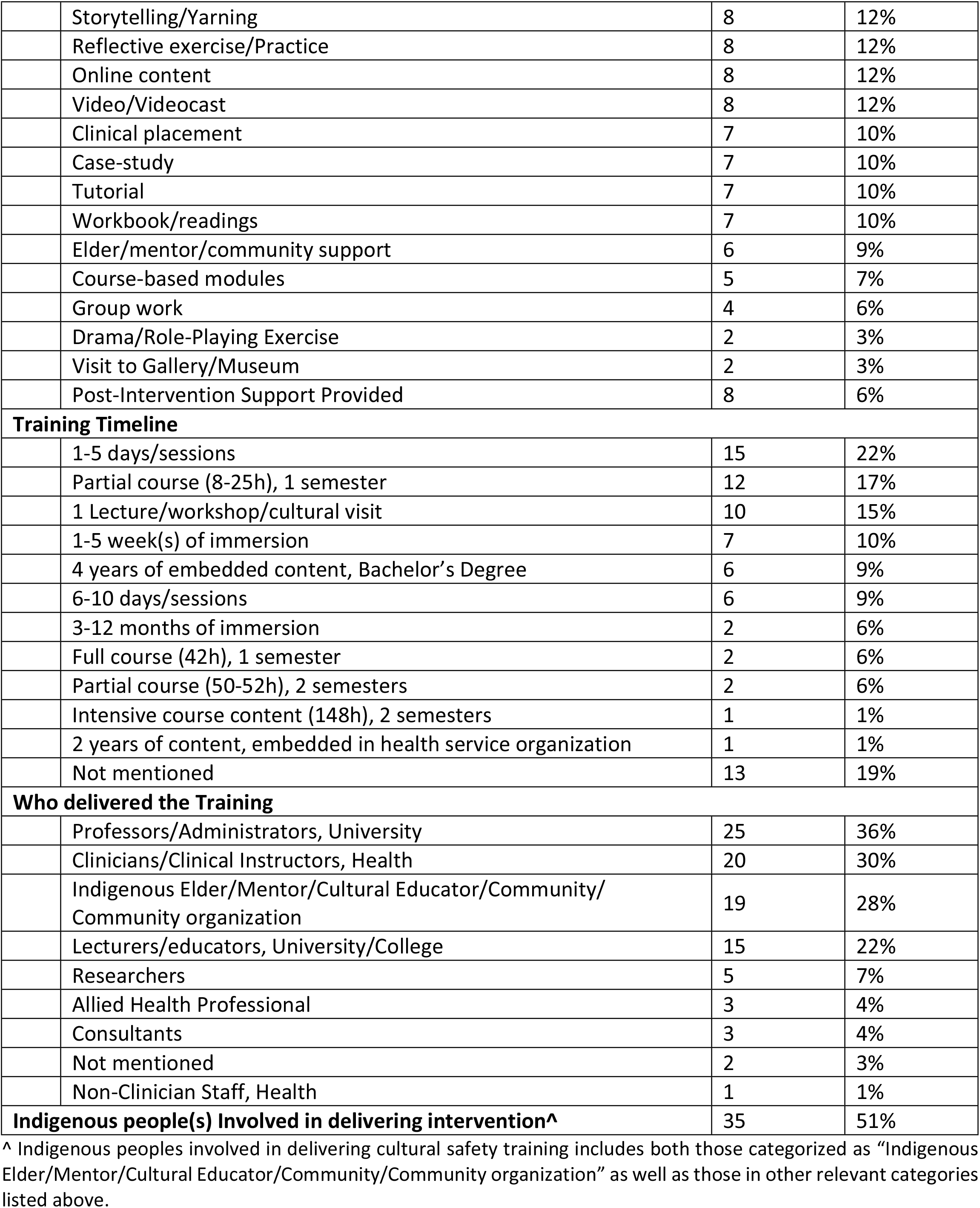
Details of Indigenous Cultural Safety Training Interventions

Following these modalities, storytelling/yarning (58, 66, 72, 80, 85, 93, 101, 113), reflective exercised/practice (77, 85, 89, 93, 101, 102, 110, 121), online content (66, 75, 83, 85, 104, 119, 121, 160), and video/videocasts (32, 46, 68, 75, 77, 89, 93, 107) were each described by an eighth of chosen studies (n=8; 12%). Also common among training modalities were clinical placements (43, 70, 114, 140, 143, 144, 153), case-studies (26, 43, 46, 47, 68, 110, 121), tutorials (39, 46, 89, 95, 104, 137, 140), and workbooks/readings (32, 34, 85, 102, 117, 135, 146), each of which were included as part of approximately a tenth of interventions (n=7; 10%). Finally, Indigenous Elder/mentors/community supports (n=6; 4%) (48, 76, 78, 85, 117, 146), course-based modules (n=5; 4%) (81, 93, 119, 123, 160), and group work (n=4; 3%) (26, 42, 57, 76) were each described by a few papers. The least common activities mentioned were drama/role-play exercises (142, 148) and a visit to a gallery/museum (32, 101), each mentioned by a couple of articles (n=2; 3%). Notably, eight papers (12%) described the provision of follow up support for learners of cultural safety beyond the primary training period (32, 34, 48, 59, 60, 123, 134, 153).

With respect to the training timeline, the largest proportion of articles detailing a cultural safety training intervention described the training as lasting 1-5 days/sessions (n=15; 22%) (26, 37, 42, 57, 60, 62, 77, 86, 103, 105, 108, 110, 134, 142, 148). Following this, in approximately one-sixth of selected articles, the training involved part of a one-semester undergraduate course (8-25 hours) (n=12; 17%) (41, 46, 49, 72, 74, 75, 80, 84, 89, 91, 104, 113). The shortest training involved a one-off lecture/workshop/cultural visit, which was mentioned in an estimated one-sixth of articles (n=10; 15%) (47, 69, 79, 81, 93, 101, 117, 124, 134, 148), followed by a 1-5 week(s) immersive experience in one of ten articles (n=7; 10%) (92, 102, 106, 114, 144, 153, 160). The longest training experience involved embedded content across a four-year bachelor’s degree (unknown total hours) (42, 43, 121, 123, 140, 146), followed by training of 6-10 days/sessions (51, 57, 77, 83, 95, 114), both occurring in one of ten articles (n=6; 9%). Three training durations, namely, 3-12 months of immersion (89, 143), a one-semester undergraduate course (42 hours) (39, 137), and a partial undergraduate course over two semesters (50-52 hours), (32, 70), similarly occurred in an estimated one-fifth of papers. The least frequent training timelines were included in only one article each (n=1; 1%), namely an intensive course over two semesters (148 hours) (34), and two years of content embedded in a health service organization (111). Nearly one in five articles did not include details on the duration of cultural safety training (n=13; 19%) (35, 36, 48, 65, 66, 68, 76, 85, 107, 123, 126, 135, 154).

In terms of delivering cultural safety training interventions, university professors/administrators were mentioned in approximately twenty-five articles that described a cultural safety training (25; 36%) (26, 32, 34, 43, 46, 57, 65, 66, 68, 72, 74, 77, 85, 86, 93, 104, 107, 108, 110, 113, 114, 117, 121, 153, 160). Clinicians/clinical instructors (health professions) (n=20; 30) (32, 37, 41, 43, 48, 57, 60, 62, 72, 77, 79, 90, 102, 106, 108, 113, 123, 124, 143, 160), and Indigenous Elders/mentors/cultural educators/community/ organizations (n=19; 28%) (48, 78, 86, 92, 93, 95, 102-104, 106, 108, 110, 113, 114, 117, 143, 144, 146, 153) were each mentioned in approximately one-third of relevant articles. University/college lecturers/educators delivered the training in an estimated one-fifth of articles (n=15; 22%) (39, 47, 49, 75, 76, 78, 81, 83, 85, 89, 104, 108, 134, 137, 142), followed by researchers in approximately one in twenty-five articles (n=5; 7%) (47, 62, 78, 93, 110). Allied health professionals (58, 140, 148) and consultants (37, 60, 111) were each mentioned in the context of training delivery in only a few articles (n=3; 4%). Two articles provided no details on who delivered the training (42, 116), while one article described non-clinician health staff as responsible for training delivery (26). Finally, half of the articles that described a cultural safety training reported that Indigenous people(s) were involved in delivering the intervention (n=35; 51%) (37, 39, 46-49, 57, 58, 60, 62, 72, 77-79, 81, 85, 86, 89, 92, 95, 101, 103, 104, 106-108, 110, 113, 117, 137, 143, 144, 146, 148, 153).

### 3.3 Evaluations of Indigenous Cultural Safety Training Interventions

Nearly one-third of papers selected for this review described an evaluation of a cultural safety training intervention (n=61; 46%) (see Table 8). These particular studies involved a broad range of objectives, with the largest proportion of papers (n=19; 31%) aimed broadly at understanding leaners’ experiences, perceptions, needs and/or preferences related to a cultural safety training in which they participated (32, 35, 41, 51, 58, 62, 70, 72, 101-104, 107, 108, 114, 123, 134, 137, 144). Another paper similarly focused on understanding perceptions of a given training intervention but from the perspective of Indigenous community members rather than participants (n=1; 2%) (32).

**Table 8:** Details of Evaluations of Indigenous Cultural Safety Training Interventions

| Evaluations of a Cultural Safety Training Intervention |  | Number<br>(n=61) | Percent<br>(%) |
| --- | --- | --- | --- |
| **Evaluation Focus/Objectives** |  |  |  |
| **Learner experiences, perceptions, needs and/or preferences** |  |  |  |
|  | Participants' general perceptions, experiences, preferences, learning needs and satisfaction with intervention | 19 | 31% |
|  | Indigenous community members' perceptions of intervention | 1 | 2% |
| **Intervention outcomes** |  |  |  |
|  | General learning experiences/outcomes/barriers | 16 | 26% |
|  | New skills/behaviours/practices | 15 | 25% |
|  | New knowledge/awareness/attitudes/confidence | 11 | 18% |
|  | Perceptions of Indigenous peoples/health issues | 1 | 2% |
|  | Learners' receptivity/resistance to intervention content | 1 | 2% |
| **Intervention and/or behaviour change processes** |  |  |  |
|  | Intervention planning and implementation process | 2 | 3% |
|  | Behaviour change processes | 2 | 3% |
| **Relationship between intervention exposure and outcomes** |  |  |  |
|  | Between learners' perceptions of Indigenous peoples and their intervention participation | 1 | 2% |
| **Intervention context** |  |  |  |
|  | Enabling factors within organizational context | 1 | 2% |
|  | Impact on organization processes and priorities | 1 | 2% |
|  | New organizational policies, positions, capacity, and mandate | 1 | 2% |
| **Methodological approaches adopted** |  |  |  |
|  | Qualitative and quantitative methods | 32 | 52% |
|  | Qualitative methods only | 20 | 33% |
|  | Quantitative methods only | 9 | 15% |

| Research methods utilized |  |  |
| --- | --- | --- |
| **1. Pre-Intervention** |  |  |
| Pre-survey | 13 | 21% |
| Pre-survey, with open-ended questions | 4 | 6% |
| Pre-focus group | 2 | 3% |
| Pre-test | 1 | 2% |
| **2. Mid-Intervention** |  |  |
| Mid-researcher observations/reflections | 4 | 6% |
| **3. Post-Intervention** |  |  |
| Post-survey | 24 | 39% |
| Post-surveys, with open-ended questions | 12 | 20% |
| Participant Interview | 11 | 18% |
| Oral/written Learner feedback | 6 | 10% |
| Post-focus group | 6 | 10% |
| Learners' reflections/case scenarios | 4 | 6% |
| Learner journal entries/digital storytelling | 3 | 5% |
| Post-test | 1 | 2% |
| Talking Circles | 1 | 2% |
| Analysis of developed curriculum | 1 | 2% |
| Post-researcher observations/reflections | 1 | 2% |
| **4. Delayed Post-Intervention** |  |  |
| Delayed post-survey (3-55 months) | 2 | 3% |
| Delayed learners' reflections (2-3 weeks) | 1 | 2% |
| **Limitations stated for chosen methods** | **33** | **54%** |

Other common evaluation approaches involved exploring participants’ reports of learning experiences/outcomes/barriers (n=16; 26%) (35, 41, 49, 76, 77, 91-93, 95, 105-108, 111, 123, 148), and the new skills/behaviours/practices that learners acquired (n=15; 25%) (51, 57, 81, 86, 99, 106, 107, 111, 113, 115, 117, 124, 133, 144, 148). Similarly common, reported in nearly one-fifth of relevant papers, involved understanding learners’ new knowledge/awareness/attitudes/confidence since the intervention (n=11; 18%) (47, 80, 81, 84, 91, 92, 105, 110, 111, 148, 153). Other evaluation approaches included learners’ perceptions about Indigenous peoples and/or their health issues (91), as well as learners’ receptivity/resistance to intervention content (89), as mentioned in one intervention evaluation each (n=1; 2%). A less common approach to evaluating cultural safety training interventions among the selected articles involved exploring processes, and specifically intervention planning and implementation processes (n=2; 3%) (34, 66), and leaners’ behaviour change processes (n=2; 3%) (113, 124).

In terms of the methodological approaches adopted to evaluate cultural safety training interventions, just over half of relevant articles describe utilizing both qualitative and quantitative methodologies (n=32; 52%) (32, 35, 46, 47, 51, 57, 60, 62, 66, 70, 72, 74, 76, 77, 81, 86, 89, 91, 93, 99, 102, 106-108, 113, 117, 123, 134, 137, 144, 148, 153). Another third of articles outlined the use of qualitative methods only (n=20; 33%) (34, 41, 43, 49, 54, 56, 58, 64, 65, 78, 80, 92, 95, 98, 101, 104, 111, 124, 133, 154), while approximately one in six papers used quantitative methods only as part of their intervention evaluations (n=9; 15%) (39, 61, 84, 105, 110, 115, 146, 156, 158). A broad range of research methods were taken up to evaluate the influence and/or impact of cultural safety training interventions on participants. Data collection methods prior to the intervention involved pre-surveys (n=13; 21%) (66, 70, 72, 81, 86, 89, 91, 93, 107, 111, 115, 123, 153), only four of which included open-ended, qualitative questions (n =4; 6%) 18, 48, 70, 76). Less common were pre-focus groups (n=2; 3%) (123, 154), and a pre-test (n=1; 2%) (110). A small number of evaluation papers undertook observations/reflections half way through the interventions (n=4; 6%) (72, 89, 92, 111).

The greatest range of data collection methods occurred immediately following the intervention, with post-surveys the most common approach, mentioned by two-thirds of relevant articles. A large proportion of these post-surveys focused only on quantitative data (n=24; 39%) (32, 35, 39, 47, 49, 66, 70, 72, 76, 84, 86, 89, 102, 103, 107, 108, 111, 114, 115, 123, 134, 137, 144, 153), while others also included open-ended qualitative questions (n=12; 20%) (49, 51, 57, 58, 62, 81, 91, 93, 95, 104, 108, 146). Participant interviews were also common following an intervention, mentioned in nearly a fifth of relevant articles (n=11; 18%) 4, 12, 51, 60, 63, 78, 82, 91, 92, 109, 134). Oral or written learner feedback (n=6; 10%) (70, 76, 77, 86, 101, 108), and post-focus groups (n=6; 10%) (32, 86, 92, 102, 123, 154) were each mentioned in a tenth of papers describing an training evaluation, followed by learner reflections or case studies (n=4; 6%) (92, 108, 124, 153) and learner journal entries or digital storytelling (n=3; 5%) (66, 72, 102), as described in a quarter of relevant papers. A Post-test (110), Talking Circles (108), Analysis of developed curriculum (34), and Post-researcher observations/reflections (107) were each mentioned in only one relevant study (n=1; 2%). Finally, a small number of relevant papers undertook data collection after a period of delay following a cultural safety training intervention. These methods included a delayed post-survey undertaken at various stages between 3-55 months following the intervention (n=2; 3%) (47, 113). While a third paper described delayed learner reflections collected at 1-2 weeks following the intervention (n=1; 2%) (92). Finally, of the papers that described a cultural safety training intervention, only approximately half of these papers presented the limitations of the methods chosen for the intervention evaluations (n=33; 54%) (34, 39, 46, 47, 49, 51, 54, 57, 65, 66, 72, 74, 77, 80, 81, 84, 91, 93, 95, 98, 99, 106, 107, 110, 111, 115, 117, 123, 133, 134, 144, 148, 156).

The review team chose to exclude or refocus some of the data themes that were introduced in the protocol paper and proposed as part of this review study. The reasons for these changes were varied. First, data related to some of the data themes were largely homogenous. These themes included, for example, the rationale for providing training on cultural safety (or related) concepts -- reasons which overwhelming involved reducing disparities between Indigenous people and relevant settler populations on health outcomes (for health-related papers) or education achievement (for education related papers). Data was also not presented in this paper on themes wherein the data was not described consistently across selected studies. This included themes such as description of training components, number of people who completed the training, and details of post-training supports provided. Similarly, data on who sponsored or developed the training intervention, as well as the duration of the training intervention and whether it was implemented in rural or urban settings were largely incomplete, with a relatively small proportion of articles including fulsome descriptions for these themes. For its part, data on ‘the aim/objective’ of the evaluation tended to be very general, including reasons such ‘to evaluate a cultural safety training intervention’, and was therefore replaced with ‘evaluation focus’, which instead centred on the *nature* of data collected (i.e. participants perceptions, intervention processes or outcomes etc.). Finally, the evaluation objectives and recommendation were not analyzed and included in this review because this data was rich in description and synthesizing the findings would require undertaking content analysis, which the authors determined were beyond the scope of this review, particularly given the time frame that was required to complete this review.

## 4. DISCUSSION

### 4.1 Summary of Evidence

This study found a significant growth in research about Indigenous cultural safety training in the fields of health, education, and social work over the last few decades, and primarily within the last ten years. The increased attention to cultural safety and related concepts has been fuelled by national initiatives to address the historical injustices against Indigenous peoples. For example, Canada’s Truth and Reconciliation Commission’s Calls to Action (2015) set out the need for cultural safety training for healthcare professionals, and federal funding for post-secondary institutions and educators to establish national research programs in collaboration with Indigenous peoples to advance Reconciliation (24). A similar 2008 initiative in Australia, entitled Closing the Gap (25), called for mutli-sectoral action in collaboration with Aboriginal communities to improve Indigenous health, which was followed in 2019 by the development of a National Indigenous Agency in 2019 responsible for leading and coordinating the implementation of the strategy (161). Australia’s commitment to action was published almost a decade before Canada’s Truth and Reconciliation Commission’s Final Report and Calls to Action, which may explain why an overwhelming majority of studies selected for this review were published in Australia – nearly 2.5 times the number of studies published in Canada, and more than three times the number published in the United States.

Only a fifth of papers selected for this review included a named Indigenous author, while an estimated one-tenth of papers failed to specify even the broad Indigenous population relevant to the study (e.g. First Nations, as in Canada). These findings suggest that greater commitment to meaningfully engage Indigenous communities on projects concerning them is urgently needed by researchers and educators working to advance cultural safety in the health, social work, and education fields (162, 163). The greatest share of articles describing a cultural safety training tended to target the field of nursing/midwifery followed by medicine/physicians. This finding is unsurprising considering that nurses/midwives tend to comprise the largest share of the health service workforce followed by physicians (164). The education field, for its part, comprised only a tenth of selected articles, suggesting that much more is needed to improve access to cultural safety training for educators. Across the four broad fields included in this study, approximately half of the population targeted by the articles were students while the other half were professionals. This is an important finding as it suggests that not only are students learning about cultural safety within their post-secondary education programs, but so are instructors, mentors and employers working in health, social work, and education fields. This suggests that professionals may have the capacity to contribute to enabling environments for new graduates to practise cultural safety when they begin their careers.

Only half of selected papers described and/or cited a cultural safety or similar concept. This is a somewhat surprising finding given that this review sought to understand the concept of cultural safety and how training interventions have attempted to facilitate its achievement. It is notable that many different concepts and definitions related to cultural safety emerged from the selected studies and were often used interchangeably. These concepts, in part, include cultural competence, cultural sensitivity, cultural responsiveness, cultural knowledge, cultural awareness, and cultural humility. Confusion related to these cultural concepts in selected studies was made worse by several factors, including: 1) few attempts to justify why a given concept was chosen over another; 2) original sources for these concepts tended to not be clearly cited; and 3) little demonstrated understanding and contextualization of the literature wherein these concepts emerged. Similarly, and perhaps unsurprisingly given the current nascent state of this literature, less than a fifth of papers provided a critique of a cultural safety or similar concept.

There is evidence to suggest there is some convergence around the geopolitical origins of two concepts, namely, cultural safety and cultural competence. Cultural safety emerged in New Zealand by a Māori nurse working within the health system at a time when the country was primarily bi-racial, comprising Indigenous peoples and Western European settlers (28). Cultural competence, for its part, is a term developed in the United States within the transcultural nursing movement and in response to nurses’ lack of understanding of the unique health needs of immigrants (128). In addition to the convergence of these two concepts, Lauren Baba, a Canadian researcher at the National Collaborating Centre for Indigenous Health in British Columbia, published a 2013 report that draws upon multiple sources to conceptualize and articulate the distinctions between cultural safety, cultural competence, and other similar concepts, including cultural awareness and cultural sensitivity (165). These four cultural related concepts and their definitions are outlined in Table 9 (Defining Cultural Safety, Cultural Competence and Similar Concepts).

**Table 9:** Defining Cultural Safety, Cultural Competence and Similar Concepts. (as set out in Baba, 2013)

| Concept | Definition |
| --- | --- |
| **Cultural Awareness** | The acknowledgement and understanding of cultural differences by focusing on the 'other' and the 'other culture'. Does not consider the political or social-economic influences on cultural difference and does not require an individual to reflect on his/her own cultural perspectives. |
| **Cultural Sensitivity** | Recognizes the need to respect cultural differences. Involves exhibiting behaviours that are considered polite and respectful by the persons of other cultures. Focuses on the 'other' and the 'other culture' and does not require an individual to reflect on his/her own cultural perspectives. |
| **Cultural Competence** | Skills and behaviours that help a practitioner provide quality care to diverse populations. While cultural competence can build upon self-awareness, it is limited by reducing culture to a set of skills for practitioners to master, and over-emphasizes cultural differences as the course of conflict between healthcare providers and diverse populations. |
| **Cultural Safety** | Cultural safety within an Indigenous context means that the educator, practitioner or professional, whether Indigenous or not, can communicate competently with a patient in that patient's social, political, linguistic, economic, and spiritual realm. Moves beyond cultural sensitivity to analyzing power imbalances, institutional discrimination, colonization, and colonial relationships as they apply to healthcare. |

Both academic research involving these cultural related concepts and training programs that seek to facilitate their development, would benefit from considering, comparing and contrasting the concepts set out by Baba (165). Not only would consolidating these terms across the literature work to further develop the scholarly work in this area, but also it would facilitate the advancement of the field more broadly by furthering the understanding of which concepts are preferable in different contexts and how to ensure their achievement through targeted training programs. This field of research would also benefit from advancing understanding on how these different concepts relate to one another in practice, how they should be applied, and the merits and drawbacks of each. Moreover, careful attention must be paid to the contexts in which these concepts are applied given the unique socio-political and historical situations in which some of them have emerged (28, 128).

Half of the papers selected for this review described a cultural safety training. While a substantial proportion of these training interventions involved Indigenous people(s) in the development processes, the specific roles they played were rarely clear. These findings highlight the need to ensure Indigenous voices are central to identifying and prioritizing the content to be included in Indigenous cultural safety trainings (162). A substantial variety of training modalities was apparent in the training interventions, with teaching or lectures, workshops, and immersive experiences or community visits among the most common. Similarly, there was a huge range in the timeline or duration of cultural safety trainings. Interventions varied from several individual sessions to one or more months of immersive experience, to fully embedded content within a post-secondary course or throughout an entire undergraduate degree program. Even the actors who delivered the training ranged significantly, from university professors and health service clinicians, to lecturers, researchers, and consultants. While approximately half of training interventions involved Indigenous Elders, experts, and other community members in the process of training delivery, the roles played by Indigenous peoples vis-à-vis other actors that were involved was not always clearly described.

There was little consistency in the objectives and methods to evaluate Indigenous cultural safety training interventions. Some evaluations focused on the experiences of learners and how they made sense of those experiences, while others looked at specific outcomes or processes (or the relationships between these two factors). There were also evaluations that considered the contexts within which the interventions were implemented and how these contexts influenced the interventions. Methodological approaches chosen for intervention evaluations also varied widely, including qualitative, quantitative, and mixed methods approaches. Most studies collected data after a cultural safety intervention, while fewer considered relevant data prior to, during, or following a delay after an intervention. Pre- and post-surveys were the most common method adopted (with some including open-ended questions), followed by interviews, focus-groups, and methods involving written reflections, case studies, journal entries and/or storytelling. Notably, only half of the studies included in this review set out the limitations of the methods chosen for intervention evaluations.

### 4.2 Limitations

There is a risk that some relevant studies may have been missed, due to selection of databases or the exclusion of grey literature. The language skills of the researchers on the team also limited the search to English publications, where French, Spanish or Indigenous languages may have been used within the selected geographic regions. Furthermore, search terms for populations did not include Indigenous nation-specific names (e.g. Mohawk, Cree), and thus publications that did not use overarching terms like Indigenous or First Nations, may have been missed.

### 4.3 Conclusions and Recommendations

The variation across studies that described a cultural safety training and/or an evaluation of a cultural safety training intervention reflects the reality of an emerging field of practice and research. It is an imperative that future work on cultural safety trainings engage Indigenous peoples at the outset and ensures their inclusion at every stage of the process, including training development, implementation, and evaluation. Moreover, to advance the field of cultural safety training in the fields of health, social work and education, research publishing rich detail is required on all aspect of developing, implementing, and evaluating interventions.

Evaluation research on cultural safety training that adopts exploratory approaches to understanding experience can identify salient aspects of training to be tested more rigorously in follow up studies. However, studies evaluating the impact of training on learning and professional practice could benefit from adopting best practices in implementation science to understand how implementation processes influence intervention outcomes. Moreover, incorporating social theory on learning (166) and behaviour change (167) into the development of cultural safety training interventions and their evaluations, can facilitate the understanding both of participants’ experience with learning about cultural safety and of how learning is applied in practice, as well as the factors that facilitate or pose barriers to these processes.

This scoping review presents the substantial work that has been undertaken over the last twenty-five years in Canada, Australia, New Zealand, and the United States to deliver Indigenous cultural safety training interventions for the purpose of addressing ongoing anti-Indigenous racism and its historical legacy in national, publicly funded health and education systems. While the research findings from this review set out the substantive progress toward achieving cultural safety through training interventions, this field of research remains in its infancy. Future research on cultural safety and related training interventions requires greater clarity in conceptualization of cultural safety and similar concepts, and more robust and context sensitive approaches considered in evaluating such interventions. Moreover, future work in this area is required to identify and how and when cultural safety and related concepts should be applied, and this is likely to vary across countries, regions, and relevant Indigenous groups.

To advance future research and practice on Indigenous cultural safety, relevant Indigenous groups must be meaningfully engaged throughout the entire duration of research and practice. A 2021 report published by the Yellowhead Institute on Canada’s progress toward Reconciliation concluded that symbolic actions were being prioritized by the Canadian Government over lasting permanent and structural changes necessary to transform the country’s relationship with Indigenous peoples (168). It’s imperative that current momentum to advance the field on Indigenous cultural safety training is capitalized upon to address anti-Indigenous racism and the legacy that colonial governments continue to impose upon Indigenous peoples in counties such as Canada.

## 4.5 Researcher Contributions

**TM:** Led data extraction and analysis processes, completed one full data extraction, analyzed a significant portion of data, and drafted and edited the manuscript.

**JQ:** Undertook a significant portion of data extraction and analysis

**LHe:** Undertook a significant portion of data extraction and some data analysis.

**AB:** Created data extraction tool, ran database searches, screened articles, contributed to methodology section, and undertook some data extraction.

**LHo**: Undertook screening of some articles and some data extraction. **PV:** Undertook screening of some articles and some data extraction. **BT:** Undertook screening of some articles and some data extraction. **ER:** Undertook some data analysis.

**EDR:** Provided guidance on screening, data extraction and data analysis and reviewed draft paper.

**AMP:** Supervised project and reviewed draft paper.

## Appendix: Image

**Image 1:**
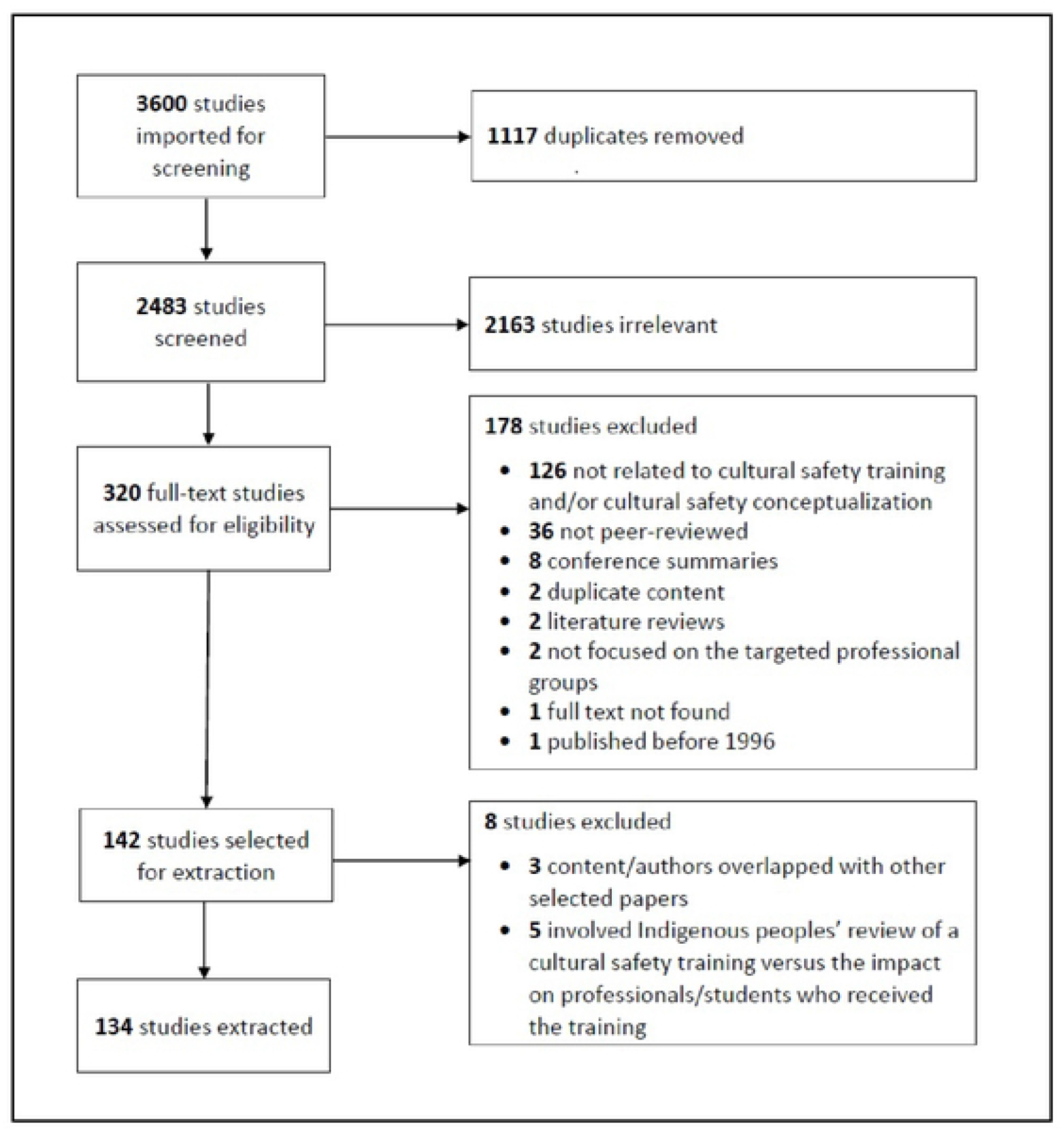
Prisma Flow Diagram

